# *Borrelia burgdorferi bb0164* encodes a Na^+^/Ca^2+^ antiporter homolog with a novel role in Mn^2+^ homeostasis and infectivity

**DOI:** 10.64898/2026.06.16.732666

**Authors:** David Gafford-Gaby, Brandee L. Stone, Vanessa M. Ante, Savana M. Green, Kennedy L. Coleman, Madeline D. Reeves, Kristin L. Rosche, Dana K. Shaw, Jenny A. Hyde

## Abstract

*Borrelia burgdorferi*, the Lyme disease causative agent, relies on trace metals for motility, growth, and virulence in the absence of metal transport homologues encoded in the genome. Previous studies characterized borrelial metal transporter (*bmtA*) as a manganese (Mn) transporter and speculated that it is the sole Mn transporter. An oxidative stress transposon library screen identified *bb0164*, annotated as a calcium/sodium antiporter, as having a putative metal binding domain. The transposon mutant lost the ability to internalize Mn suggesting *B. burgdorferi* is using a non-canonical protein for metal transport. In this study, a *bb0164* deletion and complement were generated in *B. burgdorferi* 5A4-NP1 to evaluate for trace metal transport, virulence regulation, resistance to oxidative stress, and infectivity. Our data demonstrated that the loss of *bb0164* resulted in a significant reduction in internalized Mn, increased sensitivity to oxidative stress, dysregulation of the BosR-RpoS virulence pathway, and a loss of infectivity in mice. The loss of *bb0164* resulted in elevated *rpoS*, *ospC*, and *dbpA* expression and production, while *bosR* was not altered transcriptionally or post-transcriptionally. *B. burgdorferi* grown in chelated complete BSK-II media showed a similar sensitivity to oxidative stress and virulence dysregulation as the *bb0164* mutant. These phenotypes were rescued by exogenous Mn and Zn without influencing the expression levels of *bb0164* or *bmtA*. AlphaFold models of BB0164 were structurally divergent from the canonical bacterial Mn transporter, *Bacillus subtilis* MntH and *B. burgdorferi* BmtA, but shared high similarity with a calcium/cation antiporter superfamily member. Together, this study characterized BB0164 as a second non-canonical Mn transporter in *B. burgdorferi* that is essential for mammalian pathogenesis and likely supports metal homeostasis along with *bmtA*. More broadly, *B. burgdorferi* uses unique and uncharacterized mechanisms for metal homeostasis that supports physiology and pathogenesis of the spirochete during mammalian infection.

**Author Summary:** Lyme disease, caused by *Borrelia burgdorferi*, is the most common vector-borne illness in the United States and can result in a chronic inflammatory disease. *B. burgdorferi* acquires most of the necessary nutrients, including trace metals, from the host due to its limited metabolic capacity. *B. burgdorferi* has evolved a manganese-centric metabolism in place of the iron primarily used by other bacteria. Little is understood about manganese homeostasis in *B. burgdorferi* with a single transporter, BmtA, characterized to date. Here, we describe a second manganese transporter, encoded by *bb0164*, that is essential for infection, protects against oxidative stress, and alters genetic regulation. We found that BB0164, an annotated ion antiporter, does not structurally align with conserved manganese transporters from other bacteria. Interestingly, our work suggests *bb0164* and *bmtA* are not subject to transcriptional regulation dependent on temperature or metal availability, differing from other bacterial manganese transporters. These findings indicate *B. burgdorferi* is uniquely using an antiporter protein for metal transport to support metal homeostasis, which further demonstrates the importance of manganese in borrelial pathogenesis.

## Introduction

*Borrelia burgdorferi* is the causative agent of Lyme disease, the most common vector-borne disease in the United States, and exists within an enzootic cycle between *Ixodes scapularis* ticks and mammalian or avian hosts [1–6]. *B. burgdorferi* causes a multisystemic inflammatory disease as the bacterial pathogen disseminates through the mammalian host and establishes colonization of secondary tissues [7–9]. *B. burgdorferi* encounters a wide range of stressors (e.g., nutrient, oxidative, pH) during its enzootic cycle that requires the spirochete to rapidly detect environmental changes for adaptation and survival [10–19]. *B. burgdorferi* is genomically and physiologically unique from model bacterial systems with a significant representation of annotated hypothetical proteins and domain of unknown function (DUF) proteins [20]. *B. burgdorferi* also lacks many biosynthetic pathways, relying heavily on scavenging host-derived nutrients and lipids that are readily utilized (e.g., cholesterol incorporated into the outer membrane) [21–24]. Adaptation is achieved through intensive monitoring and maintenance of cell homeostasis, including regulating intracellular concentrations of trace metals such as manganese (Mn) [18,25–29]. Metal homeostasis must be actively managed through sensing and transport, but little is known about how *B. burgdorferi* transports and utilizes trace metals [18,25–31].

In the host-pathogen trace metal arms race, *B. burgdorferi* has evolved to forgo iron (Fe) usage and instead relies on Mn as an important trace metal cofactor [29]. Mn promotes survival, virulence, and pathogenicity through mechanisms including as an enzyme cofactor, which leads to increased oxidative stress resistance and signal transduction regulation [7–16]. The risk of oxidative damage within a cell is decreased by utilizing the less reactive Mn instead of Fe as a cofactor for redox-prone enzymes (e.g., superoxide dismutase). Mn also forms non-enzymatic complexes that neutralize reactive oxygen species with small organic molecules facilitating Mn^2+^/Mn^3+^ redox cycling [32–36]. *B. burgdorferi* also depends on Mn as the cofactor for superoxide dismutase and for the optimal function of BosR along with zinc (Zn), a key transcriptional regulator of RpoS, which controls expression of required virulence factors essential for mammalian infection [18,29,37–41]. As a result, mechanisms to obtain and maintain non-toxic intracellular Mn concentrations that are vital for *B. burgdorferi* survival, virulence, and pathogenicity are yet to be fully understood.

Bacterial Mn transport occurs primarily through ATP-binding cassette transporters, Natural Resistance-Associated Macrophage Protein Nramp-type (MntH) secondary transporters, and P-type ATPases, which are not found in the borrelial genome [31,42,43]. To date, only one metal transporter has been characterized in *B. burgdorferi*, the *Borrelia* metal transport protein A (BmtA, BB0219), which transports Mn and is not homologous to previously characterized bacterial Mn transporters [25,44]. The ability of BmtA to transport metals and influence virulence determinants has been investigated; however, an additional putative Mn transporter has been identified in *B. burgdorferi*, BB0164 [18,25,45]. The presence of multiple, non-redundant Mn transporters in *B. burgdorferi* suggests Mn homeostasis is more complex than previously appreciated, but how these systems contribute to survival and infectivity remains unclear. BB0164 is a putative inner membrane protein that, like BmtA, is not homologous to characterized bacterial Mn transporters [45]. A *bb0164* transposon (Tn::*bb0164*) mutant showed decreased intracellular Mn suggesting transport activity [45]. BB0164 is annotated as a K^+^-dependent Na^+^/Ca^2+^ exchange protein, a family within the calcium:cation antiporter (CaCA) family and Cation Diffusion Facilitator (CDF) superfamily [45]. Bacterial and archaeal CaCA family homologues canonically transport calcium, sodium, and potassium [46–55]. To our knowledge, none have been shown to directly transport Mn, though two showed inhibition of calcium transport in the presence of excess Mn [48,54]. In contrast, CaCA homologues in plants do directly transport Mn [56–60].

Given the absence of characterized metal transport systems beyond BmtA and the atypical features of BB0164, we sought to determine whether BB0164 contributes to borrelial metal transport and borrelial virulence. We show that BB0164 functions as a second, non-redundant Mn transporter required for oxidative stress resistance, proper regulation of virulence factors under mammalian-like conditions, and to establish infection in the experimental model. These findings identify BB0164 as an essential component of Mn homeostasis and demonstrate that unconventional transport systems contribute to the unique metal biology required for *B. burgdorferi* survival and pathogenesis.

## Results

### BB0164 is structurally similar to Na^+^/Ca^2+^ exchange family transporters and structurally distinct from other bacterial Mn transporters

When the *B. burgdorferi* genome was sequenced, *bb0164* was annotated as encoding a K^+^-dependent Na^+^/Ca^2+^ exchanger like-protein [20]. Previous work screening the *B. burgdorferi* Tn library identified *bb0164* as contributing to oxidative stress survival and modulating intracellular Mn levels [45]. To determine whether the predicted structure of BB0164 is similar to other characterized Mn transporters, we modeled BB0164, *B. burgdorferi* borrelial metal transporter (BmtA), *Bacillus subtilis* Mn transporter (MntH), and *Methanococcus jannaschii* Na^+^/Ca^2+^ exchange protein (NCX_Mj) in apo and cation-bound states using AlphaFold 3 (Fig 1) [51–53,55,61,62]. Aligning these structures revealed a dissimilarity of BB0164 to BmtA or MntH but structural conservation with NCX_Mj despite sharing only 32.2% amino acid identity (Fig 1A). These relationships are further confirmed when calculating backbone Cα root mean square deviation (RMSD) values. BB0164, compared to BmtA and MntH, produce RMSD values greater than 10 Angstroms (Å) representing the distinct differences between these proteins, while BB0164 and NCX_Mj have an RMSD value of 2.969 Å indicating that the polypeptide backbone and thus overall protein structure aligns much more closely with NCX-Mj. Further, in our model E55 and E234 in BB0164 are predicted to comprise the metal-coordinating residues with contributions from backbone carbonyl oxygens from T51 and T230 and these are conserved with the ion coordinating residues E54, E213, T50, and T209 in NCX_Mj (Fig 1A). CheckMyMetal (CMM) analyses of Mn^2+^-bound BB0164 showed borderline valence (1.5) and dubious nVECSUM (0.25) scores but an acceptable gRMSD (3.6°) score. Predicted geometry was tetrahedral (borderline score), which is uncommon but not impossible to find across known Mn transporters [43,63–66]. The glutamic acid and threonine residues predicted and identified for BB0164 and NCX_Mj are distinct from those predicted to form the metal coordinating site in BmtA and MntH (Fig 1B). iPTM for Mn^2+^-bound BB0164 was 0.93 and pLDDT values for the residues predicted to coordinate Mn^2+^ in BB0164 range from 89.8 to 95.4 further suggesting BB0164 coordinates Mn (Fig S1). Finally, mapping RMSD values between apo and Mn^2+^-bound BB0164 identified regions predicted to undergo conformational changes upon coordinating Mn^2+^, specifically in the N-terminal domain (Fig 1C and S2). This is notable as NCX_Mj exhibits a similar conformational change in response to ion binding, which contributes to its exchange function [53]. Taken together, these data show the predicted structure of BB0164 is unlike established canonical bacterial Mn^2+^ transporters, including the borrelial BmtA, but is similar to the Calcium/cation (CaCA) family NCX_Mj.

**Fig 1.**
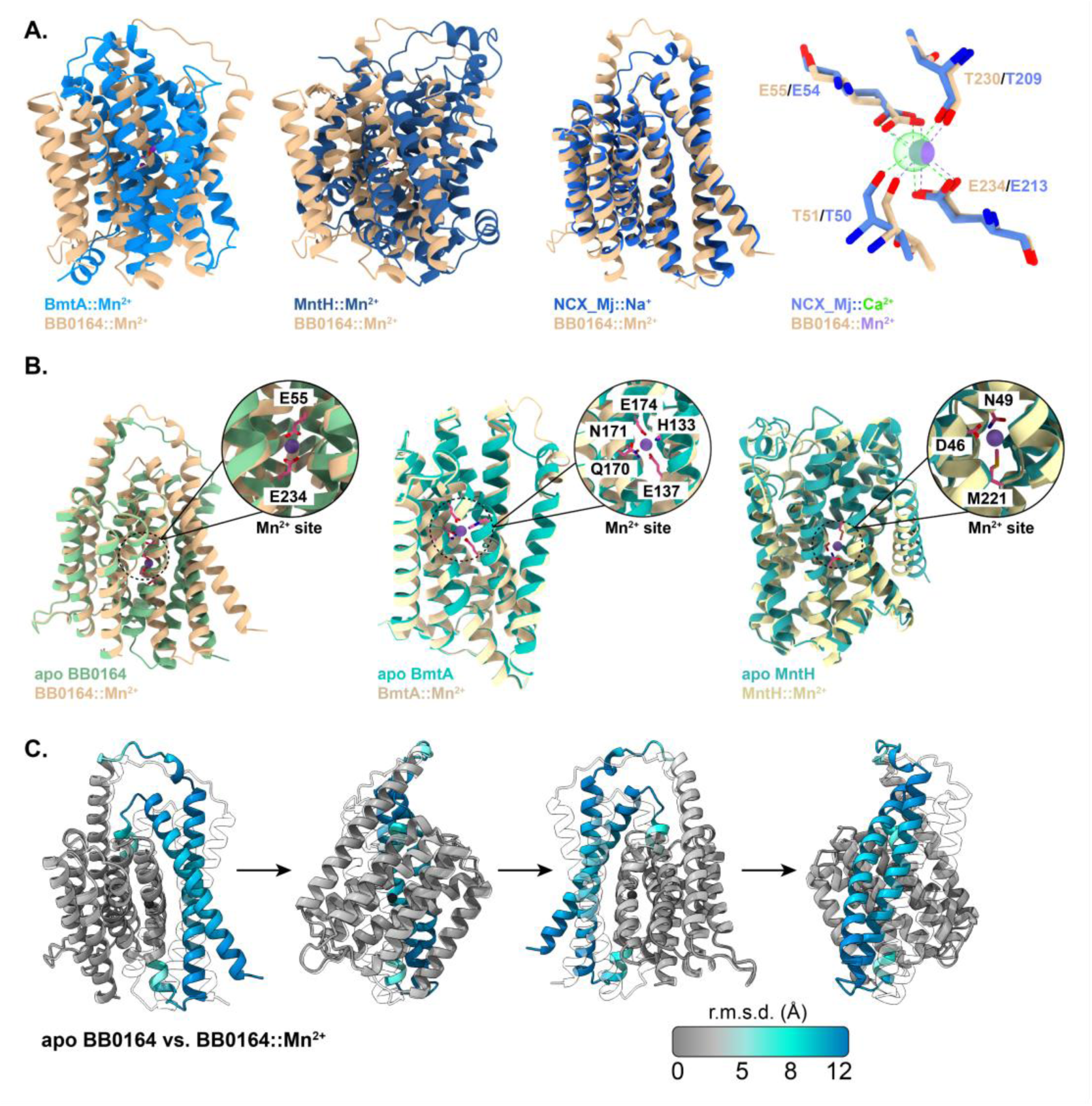
BB0164 is predicted to be structurally similar to NCX transporters. **A.** Models of BB0164 aligned with models for BmtA, MntH, and NCX_Mj. Models were generated using AlphaFold 3 and additional X-ray crystallography data (PDB ID: 5HWY and 5HXR) was used for NCX_Mj. BB0164 (tan), BmtA (blue), and MntH (blue) were modeled with Mn^2+^ (purple) and NCX_Mj (blue) were modeled with Ca^2+^ (green). **B.** Apo vs Mn-bound alignments showing key residues predicted to be involved in Mn coordination. **C.** Conformational changes predicted to occur when BB0164 coordinates Mn^2+^. Alignments and RMSD calculations were performed with ChimeraX v1.10.

### *bb0164* modulates virulence factors and is required for mammalian infectivity

Given the predicted structural homology of BB0164, we set out to characterize the function of *bb0164* by generating a targeted gene deletion and chromosomal complement of *bb0164* in wild type (WT) *B. burgdorferi* 5A4-NP1 (Fig 2A). Due to the close proximity and directionality of *bb0164* with upstream genes *bb0166* (previously identified as encoding a MalQ homolog [67,68]) and *bb0165* and with downstream gene *bb0163*, these genes are potentially organized in an operon. By RT-PCR, we confirmed co-transcription of the genes by amplifying fragments spanning the individual intergenic regions within the *bb0166-bb0163* region (Fig S3). Larger transcripts spanning *bb0166-bb0164* and *bb000165-bb0163* were also amplified across this region of interest. These findings informed our deletion and complementation strategy. When generating the *bb0164* deletion strain (*bb0164^-^*) in 5A4-NP1, *bb0164* was replaced by a *P_flgB_*-driven streptomycin resistance cassette followed by the promoter region of *bb0166* upstream of *bb0163* to prevent a polar deletion [69]. We then genetically complemented *bb0164* by inserting *P_bb0166_*-*bb0164* and a *P_flgB_*-driven gentamycin resistance cassette into an established site in the chromosomal intergenic space between *bb0445* and *bb0446* by allelic exchange [70]. All borrelial strains were evaluated for endogenous plasmid content (data not shown) and by qRT-PCR to demonstrate abrogation of *bb0164* transcription in *bb0164*^-^ and restoration of WT-level expression in *bb0164^-/+^* (Fig 2B). Furthermore, we confirmed that *bb0163* expression levels were not significantly altered in *bb0164^-^* or *bb0164^-/+^* relative to WT. Upstream genes *bb0166* and *bb0165* expression increased slightly by 1.76- and 2.06- fold, respectively, relative to WT. However, as these changes are minimal and occurring in genes upstream of the *bb0164* deletion site, they are likely not biologically significant.

**Fig 2.**
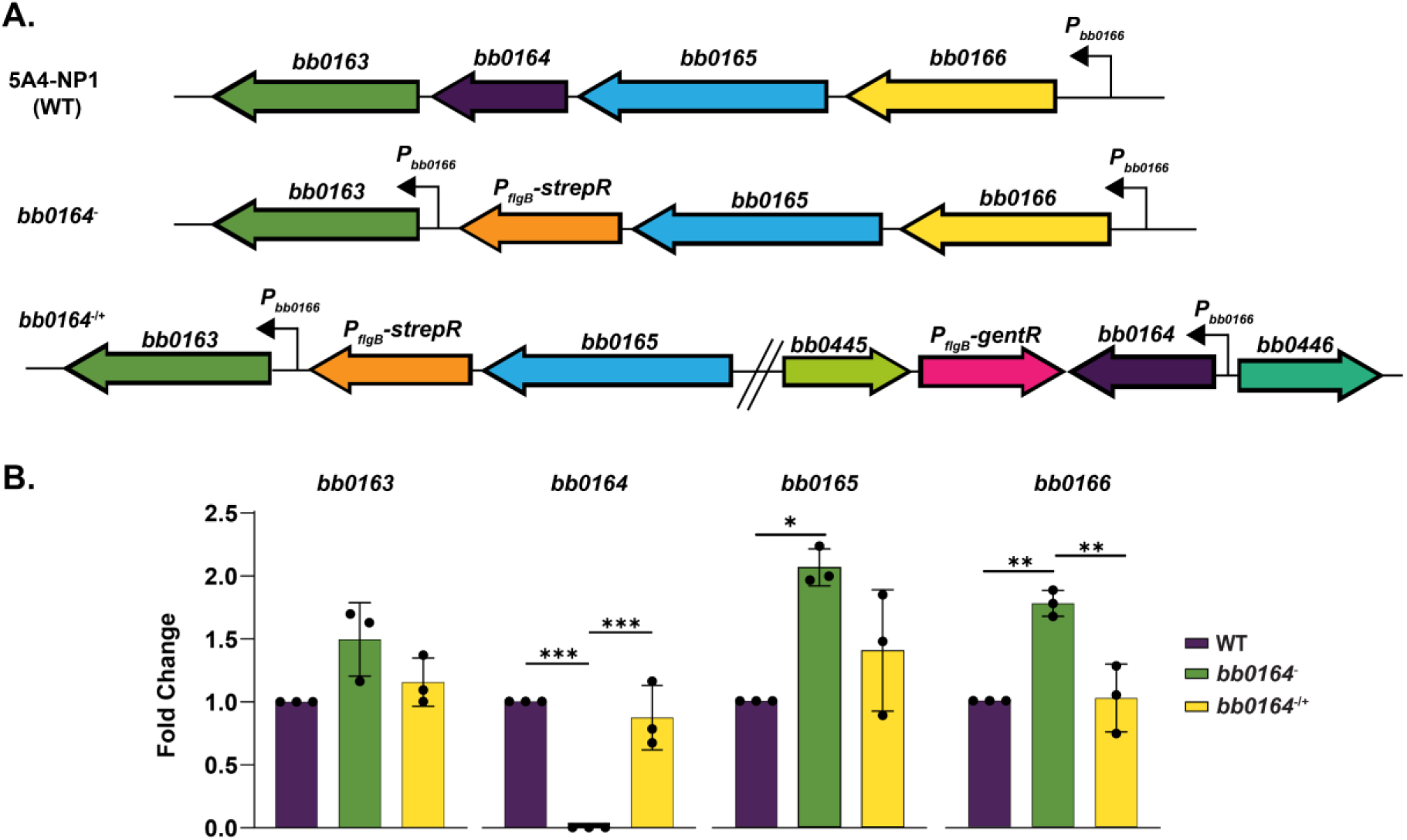
Strains used in this study confirmed by qRT-PCR. **A.** Schematic representation of the *bb0163-bb0166* in *B. burgdorferi* 5A4-NP1 (WT), *bb0164^-^*, and *bb0164^-/+^*in the intergenic chromosomal space between *bb0445* and *bb0446.* **B.** qRT-PCR analysis of operon genes in WT, *bb0164^-^,* and *bb0164^-/+^.* Data represents three biological replicates. Fold change normalized to WT for each respective gene. * p<0.05. ** p<0.01. *** p<0.001.

The borrelial mammalian-phase virulence factor regulator BosR and the genes in its downstream regulon as part of the BosR-RpoS regulatory cascade are known to be responsive to divalent metal levels [18,27], and prior work and our structural analysis suggests that BB0164 may be involved in Mn transport [45]. We therefore sought to determine if deletion of *bb0164* affected transcription of *bosR, rpoS*, and the mammalian-phase virulence factors *ospC* and *dbpA*, as well as the protein levels of some of these targets (Fig 3). WT, *bb0164^-^*, and *bb0164^-/+^*were grown at mammalian-like conditions (37°C, 5% CO_2_) and tick-like conditions (23°C, atmospheric CO_2_) [17,39,71–74]. We confirmed an induction of *bosR, rpoS, ospC,* and *dbpA* at 37 °C, 5% CO_2_ relative to 23 °C in WT *B. burgdorferi*, as expected (Fig 3A). Compared to WT*, bb0164^-^ bosR* levels were not significantly different at either 23 °C or 37 °C with 5% CO_2_, indicating that deletion of *bb0164* does not transcriptionally alter *bosR* (Fig 3A). At 37 °C with 5% CO_2_, *rpoS* levels are slightly albeit significantly elevated in *bb0164^-^*when compared to WT, with a 1.5-fold increase in *rpoS*. As expected with elevated *rpoS* levels, transcript levels of *ospC* and *dbpA* in *bb0164^-^* were elevated approximately 8-fold and 5-fold, respectively, higher than WT at 37 °C, 5% CO_2_ pointing to an overexpression of these RpoS regulon members in *bb0164^-^* (Fig 3A). Protein production was also evaluated under mammalian-like and tick-like conditions using FlaB as loading control. *bb0164^-^* displayed higher levels of RpoS and OspC at 37 °C with 5% CO_2_, but unchanged levels of BosR compared to WT (Fig 3B). These findings suggest that in *bb0164^-^,* elevated *rpoS* expression and RpoS production are occurring in the absence of BB0164 activity.

**Fig 3.**
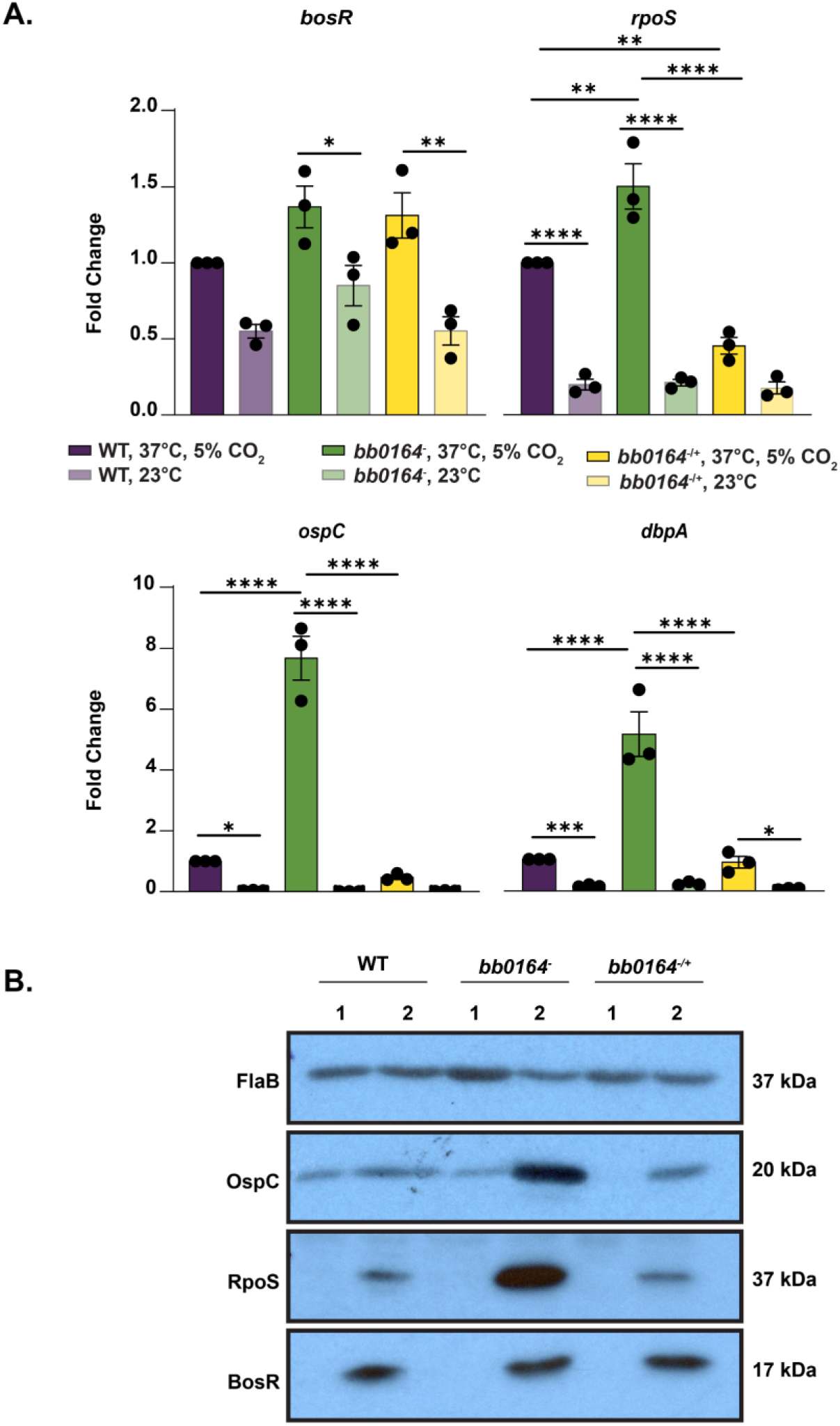
*bb0164* transcriptionally alters *rpoS* and its downstream regulon without impacting BosR production. **A.** qRT-PCR analysis of relative expression of *bosR*, *rpoS, ospC,* and *dbpA* in WT, *bb0164^-^,* and *bb0164^-/+^* grown at 37°C, 5% CO_2_ (dark shaded bars) and 23°C, atmospheric CO_2_ (light shaded bars). Fold change normalized to WT grown at 37°C, 5% CO_2_ for each respective gene. Data represents three biological and technical replicates. * p<0.05. ** p<0.01. *** p<0.001. **** p<0.0001. **B.** Western immunoblotting of WT, *bb0164^-^,* and *bb0164^-/+^* grown at 23°C, atmospheric CO_2_ (1) and 37°C, 5% CO_2_ (2) probing for OspC, RpoS, and BosR, with FlaB as a loading control. Western blots are representative of three biological replicates.

As *bb0164^-^* displayed a dysregulation of several virulence factors required for *B. burgdorferi* to establish an infection in mammals, we predicted that *bb0164* may also be required for mammalian infectivity. To investigate this, groups of C3H/HeN mice were infected by needle-inoculation with a 10^5^ dose of either WT, *bb0164^-^*, or *bb0164^-/+^* and tissues collected at 14 or 28 days post-inoculation (Fig 4). No viable spirochetes were cultured from mice infected with *bb0164^-^*, while 100% of tissues from mice infected with WT or *bb0164^-/+^* were culture positive. Such results indicate that *bb0164* is indispensable for mammalian infectivity by needle inoculation.

**Fig 4.**
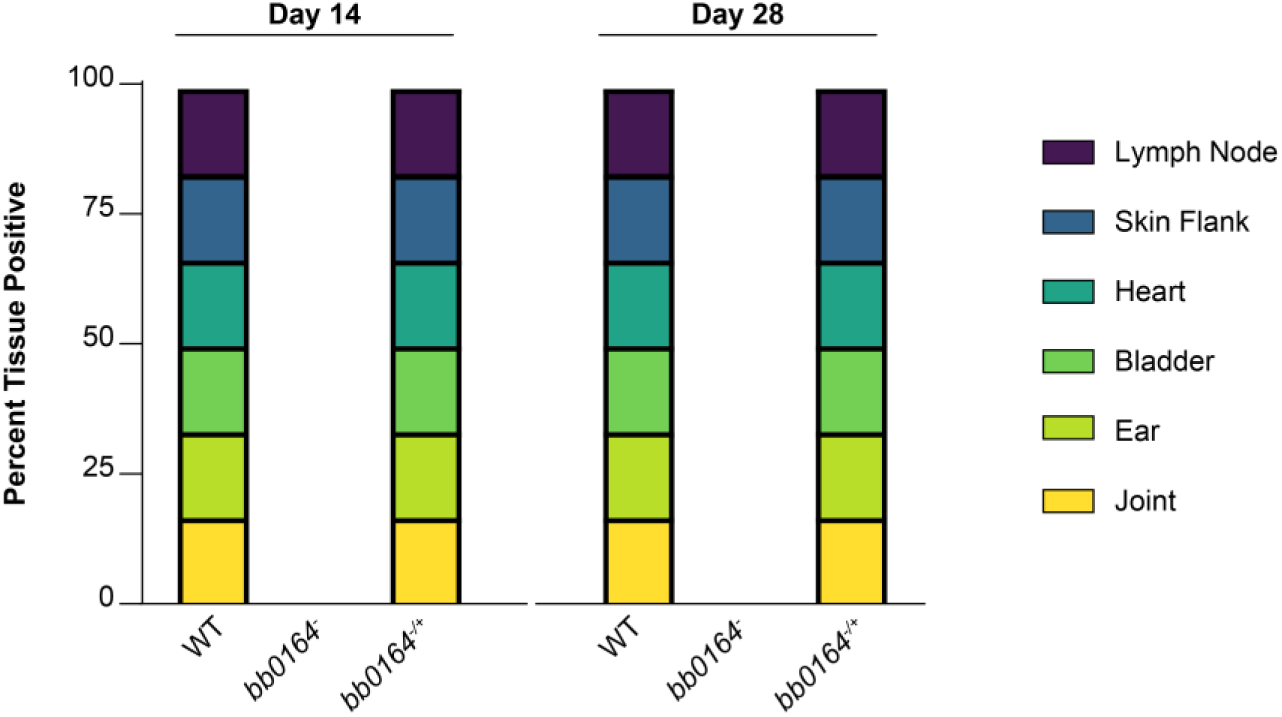
*bb0164* is required for mammalian infectivity by needle inoculation. Tissues were collected from C3H/HeN mice infected by intradermal inoculation with 10^5^ WT, *bb0164^-^,* and *bb0164^-/+^*at 14 and 28 days post-infection for outgrowth in complete BSK-II. Results represent percentage of tissues positive for *B. burgdorferi* via darkfield microscopy within 1 month.

### *bb0164* modulates intracellular Mn levels and provides protection from oxidative stress

Previous work with Tn*::bb0164* suggested a Mn transport function for BB0164 [45]. To validate this Mn transport function and investigate transport of additional trace metals and ions, inductively coupled plasma mass spectrometry (ICP-MS) was performed on WT, *bb0164*^-^, and *bb0164^-/+^* borrelial cells (Fig 5A) [18,25,26,45,75–78]. In *bb0164^-^*, we observed a 10-fold decrease in intracellular Mn relative to WT with no change in either Zn or Mg. *bb0164^-/+^* was able to partially restore Mn accumulation 7.8-fold times higher than in *bb0164^-^*, albeit to levels still 1.3-fold lower than in WT. Na accumulation was also measured given the annotation of *bb0164* as an NCX transporter; attempts to measure intracellular Ca levels by ICP-MS were unsuccessful due to the high spectral interference with Ca from the Ar^40^ gas used in the ICP-MS process. Aside from these divalent metals, *bb0164^-^*displayed a slight, though significant increase in Na ions at 1.4-fold higher than in WT that was restored to WT-levels in *bb0164^-/+^*. These data indicate BB0164 contributes to Mn accumulation into *B. burgdorferi* cells. Further, the increased accumulation of Na in *bb0164^-^*suggests that BB0164 may also contribute to modulation of Na levels in the cell, consistent with its structural homology with a Na exchanger.

**Fig 5.**
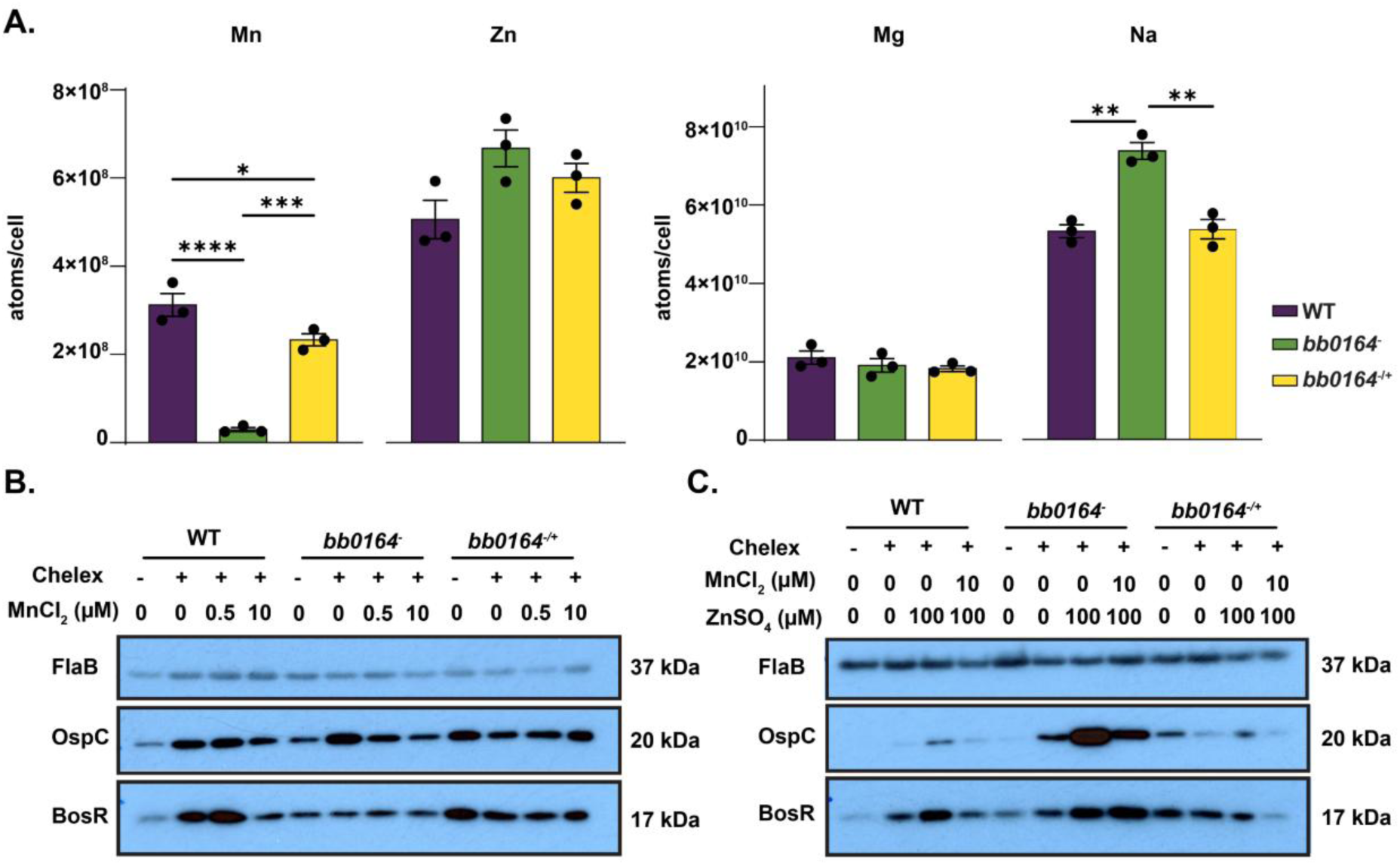
*bb0164* impacts Mn^2+^ accumulation within *B. burgdorferi.* **A.** Inductively coupled plasma mass spectrometry (ICP-MS) measurement of intracellular Mn, Zn, Mg, and Na levels in WT, *bb0164^-^,* and *bb0164^-/+^* cells grown to mid-exponential phase, with metal content normalized by post-wash cell number (∼5x10^9^ cells total). Data represents three biological replicates. * p<0.05. ** p<0.01. *** p<0.001. **** p<0.0001. **B.** Western immunoblot analysis of production of BosR, and OspC in WT, *bb0164^-^,* and *bb0164^-/+^* cells grown in complete BSK-II, chelated BSK-II, and chelated BSK-II supplemented with either 0.5 μM MnCl_2_ or 10 μM MnCl_2_. **C.** Western immunoblot analysis of BosR, and OspC in WT, *bb0164^-^,* and *bb0164^-/+^* cells grown in complete BSK-II, chelated BSK-II, and chelated BSK-II supplemented with either 100 μM ZnSO_4_ or a combination of 10 μM MnCl_2_ and 100 μM ZnSO_4_. Western blots are representative of three biological replicates.

Relative concentrations of intracellular divalent metals, particularly Mn and Zn, are known to impact borrelial virulence factors essential for adaptation to the mammalian host [18]. Specifically, Mn is known to post-transcriptionally repress the mammalian virulence factor regulator BosR, which in turn leads to reduced transcription of the alternative sigma factor *rpoS* and other elements of its downstream regulon, including *ospC* [18]. This repression by Mn can be overcome by high levels of Zn [18]. Since our data show that BB0164 modulates intracellular Mn levels, we sought to determine the effects of altered Mn and Zn availability on borrelial virulence determinants. Borrelial strains were grown in complete BSK-II and chelated complete BSK-II with and without MnCl_2_ and/or ZnSO_4_ (Fig 5B, 5C, & S4). The reduction of divalent metals in chelated BSK-II in comparison to complete BSK-II was confirmed by ICP-MS, which demonstrated a significant decrease in Mn, Zn, Fe, and Mg, but no change in Na, as expected (Fig S4A). With this media, we then assessed the ability of WT 5A4-NP1 to grow under chelated and metal supplementation at 32°C and 1% CO_2_ (Fig S4B). WT in complete and chelated BSK-II had a doubling time of 16.33 h and 23 h, respectively. WT in chelated BSK-II reached a maximum cell density of approximately 6.8x10^7^ cells/mL, nearly 3-fold lower than complete BSK-II. Chelation reduced motility but resulted in no change in cell morphology. Supplementation with 10 µM MnCl_2_, 100 µM ZnSO_4_, or both restored the doubling rate and maximum density to WT levels though cells had a prolonged lag phase. These results indicate that metal availability, particularly Zn, may play a role in the optimal *in vitro* growth of *B. burgdorferi*.

With our media chelation scheme validated, we then sought to probe our borrelial strains grown in differing Mn and Zn availabilities for BosR and OspC at an early mid-log growth phase. Growth of WT in chelated BSK-II resulted in increased BosR and OspC production relative to complete BSK-II (Fig 5B), similar to prior work that observed more OspC in chelated BSK-II in *B. burgdorferi* strain 297 [18]. Supplementation with 0.5 µM MnCl_2_ did not have a major impact on BosR or OspC levels in WT, while addition with 10 µM MnCl_2_ resulted in an expected repression of both BosR and OspC in WT. BosR levels did not change in *bb0164^-^* across these conditions. OspC increased in chelated grown *bb0164^-^* relative to complete BSKII and 10 µM MnCl_2_ supplementation. *bb0164^-/+^* had elevated BosR and OspC regardless of chelation or MnCl_2_ concentration. Collectively, these results support the notion that BosR and OspC are responsive to Mn availability which can be attributed to BB0164, in part. When 100 µM ZnSO_4_ was added to chelated BSK-II, BosR and OspC levels increased in WT, as previously reported (Fig 5C) [18]. Supplementation with 10 µM MnCl_2_ and 100 µM ZnSO_4_ displayed an intermediate phenotype in WT *B. burgdorferi*, with the presence of Zn able to overcome the repressing action of Mn on BosR and OspC. The addition of ZnSO_4_ alone and in combination with MnCl_2_ to *bb0164^-^* resulted in a substantial increase in BosR and OspC protein production when compared to WT. *bb0164^-/+^* largely restored BosR and OspC to WT-levels, with ZnSO_4_ supplementation increasing both BosR and OspC, and supplementation with both ZnSO_4_ and MnCl_2_ displaying a moderation of the repressing activity of Mn. The relative ratios of Mn and Zn have an impact on BosR and the RpoS regulon, indicating that BB0164 plays a role in influencing this ratio dependent upon metal availability.

Prior work with Tn::*bb0164* in a pooled study suggested that *bb0164* also plays a role in protecting *B. burgdorferi* from H_2_O_2_-mediated oxidative stress [45]. Likewise, previous studies with another established *B. burgdorferi* Mn transport gene, *bmtA,* found that deletion of *bmtA* rendered *B. burgdorferi* significantly more sensitive to tert-butyl hydroperoxide (TBHP)-mediated oxidative stress, suggesting that Mn transport plays a role in reactive oxygen species (ROS) protection [25]. To further investigate the contribution of *bb0164* in protecting *B. burgdorferi* from reactive species it may encounter in its enzootic cycle, we treated our WT, *bb0164^-^,* and *bb0164^- /+^* strains with H_2_O_2_, TBHP, or diethylamine/nitric oxide (DEA/NO) to evaluate borrelial survival (Fig 6). When treated with both 125 and 185 μM of H_2_O_2_, *bb0164^-^* displayed a significantly lower survival ratio with an approximate 2-fold and 1.75-fold reduction in relative outgrowth compared to WT, respectively (Fig 6A). We next sought to evaluate the impact of TBHP, an ROS stressor that primarily causes damage in *Borrelia* via lipid peroxidation [11]. Unlike with H_2_O_2_, we did not see a significant difference in *bb0164^-^* outgrowth relative to WT with either 1.5 or 2.5 mM TBHP treatment (Fig 6B). Similarly, when treated with either 1.25 or 2.5 mM of DEA/NO, a NO donor that acts as a reactive nitrogen species (RNS) stressor, *bb0164^-^* outgrowth was not differentially impacted compared to WT (Fig 6C). Combined, these results indicate that *bb0164* contributes to protection of *B. burgdorferi* from H_2_O_2_-mediated ROS specifically, but not TBHP-mediated ROS or RNS stress.

**Fig 6.**
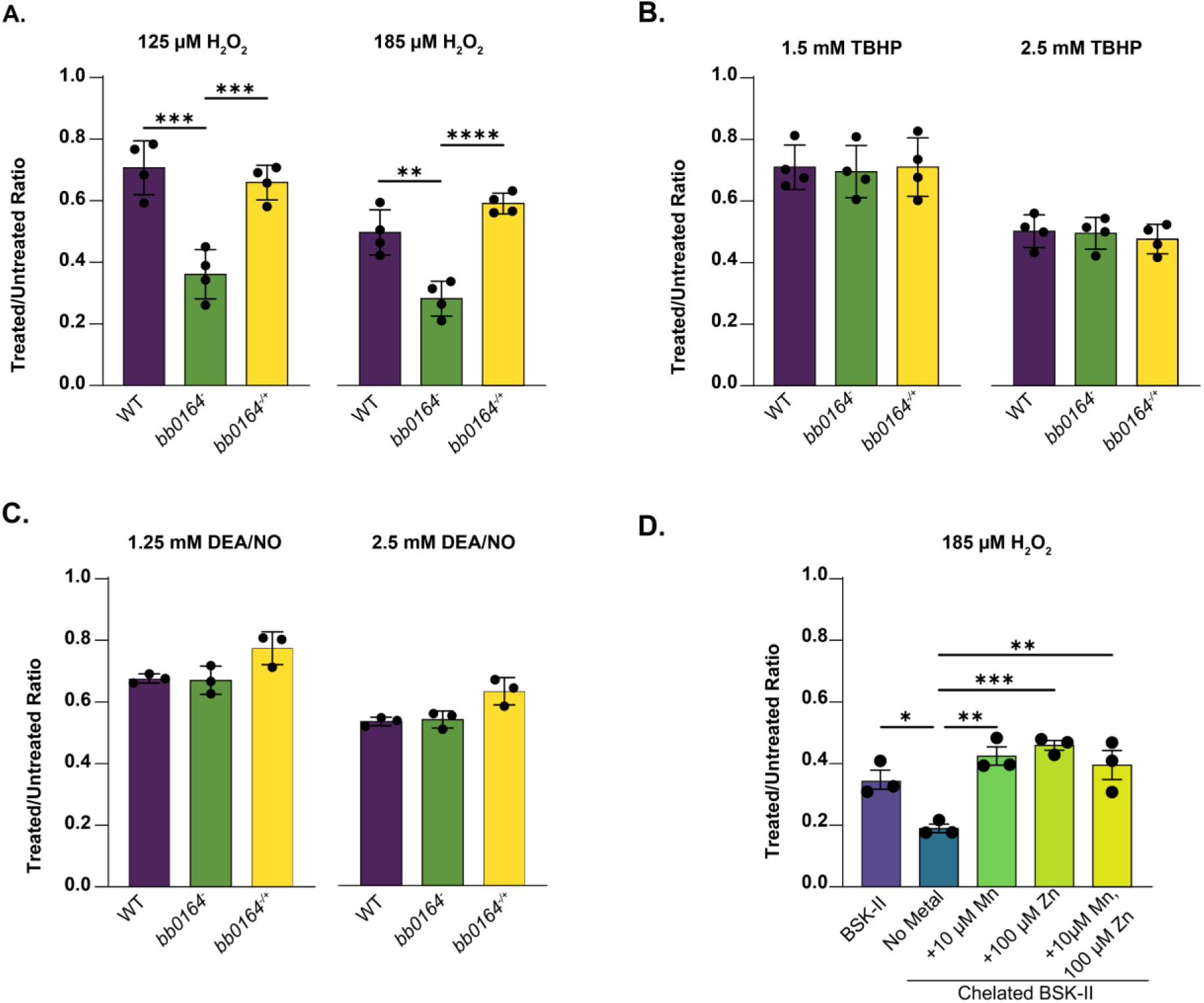
Mn accumulation by *bb0164* protects *B. burgdorferi* from H_2_O_2_ mediated oxidative stress killing. WT, *bb0164^-^,* and *bb0164^-/+^* treated with **A.** H_2_O_2_, **B.** TBHP, and **C.** DEA/NO in modified BSK-II for 2 - 4 hours and cultivated for outgrowth in complete BSK-II. **D.** WT 5A4-NP1 was grown in either complete BSK-II, chelated BSK-II, or chelated BSK-II supplemented with 10 µM MnCl_2_, 100 µM ZnSO_4_, or a combination of 10 µM MnCl_2_ and 100 µM ZnSO_4_. Samples were treated with 185 µM H_2_O_2_ for 4 hours in modified BSK-II, chelated modified BSK-II, or chelated modified BSKII supplemented with metals as appropriate. Survival was determined by cell counts of treated compared to untreated dilutions. Data represents three to four biological replicates. * p<0.05. ** p<0.01. *** p<0.001. **** p<0.0001.

Since *bb0164* plays an apparent role in intracellular Mn accumulation and H_2_O_2_-mediated ROS survival, and Mn is a known cofactor for the borrelial encoded superoxide dismutase [79,80], we hypothesized that Mn availability impacts the ability of *B. burgdorferi* to respond to ROS pressures. To test this, WT *B. burgdorferi* grown in complete BSK-II, chelated BSK-II, or chelated BSK-II supplemented with MnCl_2_ and/or ZnSO_4_ were treated with 185 μM H_2_O_2_ to assess outgrowth survival. WT grown in chelated BSK-II showed an approximate 1.8-fold reduction in outgrowth survival ratio compared to WT *B. burgdorferi* grown in complete BSK-II (Fig 6D). Strikingly, while WT *B. burgdorferi* grown in chelated media with 10 μM Mn did restore the outgrowth survival to levels seen in complete BSK-II, supplementation with 100 μM ZnSO_4_ had an identical effect. Equally surprisingly, *B. burgdorferi* grown with both 10 μM MnCl_2_ and 100 μM ZnSO_4_ showed no apparent cumulative effect and displayed a similar outgrowth ratio to cells grown in complete BSK-II. This suggests borrelial survival during H_2_O_2_ stress is broadly influenced by the availability of divalent trace metals and may not be Mn-specific.

### Expression of borrelial Mn transporters are not differentially regulated by environmental signals or metal availability

Our results suggest that BB0164 functions as a Mn importer, a function that has also previously been ascribed to the Borrelial metal transporter A (BmtA) [25]. Given the presence of two borrelial Mn transporters, we hypothesized that *bb0164* and *bmtA* may influence the regulation of the other dependent on Mn transport activity and/or under different conditions. Given this, we first sought to evaluate the effects of temperature and CO_2_ on *bb0164* and *bmtA* expression in WT, *bb0164^-^,* and *bb0164^-/+^* grown to mid-exponential phase, a more metabolically active growth phase (Fig 7A) [25,45,81]. WT grown at 23°C relative to 37 °C, 5% CO_2_ had a 1.6-fold reduction in *bb0164* expression that was statistically significant but likely not biologically significant. The deletion of *bb0164* did not alter transcriptional levels of *bmtA* at either evaluated *in vitro* condition. Interestingly, expression of *bmtA* in WT *B. burgdorferi* was not significantly altered between 23 °C and 37 °C with 5% CO_2_ at mid-exponential phase, contrasting with previous findings at stationary phase in *B. burgdorferi*, strain 297 [18]. Troxell et al. previously found that both Mn levels and *bmtA* transcript are responsive to temperature shifts in stationary phase *B. burgdorferi*, strain 297, with intracellular Mn increasing nearly 5-fold at 25°C compared to 37°C, and *bmtA* expression increasing 3.5-fold at 23°C compared to 37°C [18]. Together, our data suggests that during metabolically active growth phase, *bb0164* and *bmtA* are subject to minimal temperature- or CO_2_-dependent transcriptional regulation.

**Fig 7.**
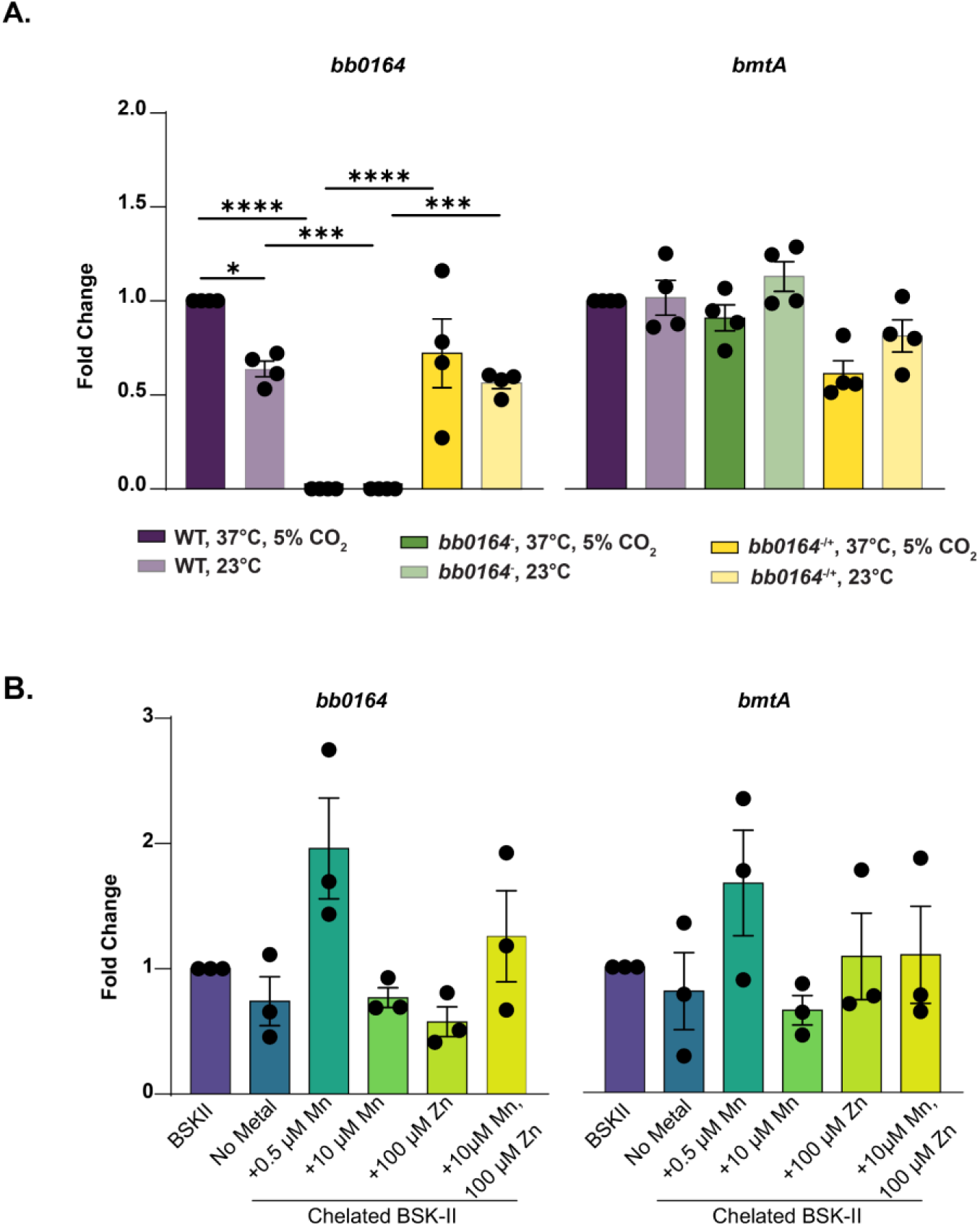
*bb0164* and *bmtA* are not transcriptionally regulated by temperature, CO_2_, or metal availability *in vitro*. **A.** qRT-PCR analysis of *bb0164* and *bmtA* relative expression normalized by *flaB* expression in WT, *bb0164^-^,* and *bb0164^-/+^* cells grown at 37°C, 5% CO_2_ (dark shaded bars) and 23°C, atmospheric CO_2_ (light shaded bars). **B.** qRT-PCR analysis of *bb0164* and *bmtA* relative expression normalized by *flaB* expression in WT cells grown in normal BSK-II, chelated BSK-II, and chelated BSK-II supplemented with 0.5 μM MnCl_2_, 10 μM MnCl_2_, 100 μM ZnSO_4_, or a combination of 10 μM MnCl_2_ and 100 μM ZnSO_4_ at 32°C, 1% CO_2_. Data represents three to four biological replicates. * p<0.05. *** p<0.001. **** p<0.0001.

Though *bb0164* and *bmtA* did not appear strongly transcriptionally responsive to temperature or CO_2_, Mn importers in other systems can be transcriptionally regulated as a result of metal availability [82,83]. To determine if metal availability in growth media alters *bb0164* and *bmtA* expression, WT *B. burgdorferi* were grown in complete BSK-II, chelated BSK-II, and chelated BSK-II supplemented with 0.5 µM MnCl_2,_ 10 µM MnCl_2_, 100 µM ZnSO_4_, or 10 µM MnCl_2_ and 100 µM ZnSO_4_ (Fig 7B). Expression of *bb0164* and *bmtA* were not significantly different under any metal availability condition, showing that neither are transcriptionally regulated by Mn or Zn availability under our *in vitro* conditions. Collectively, our data indicate that neither *bb0164* nor *bmtA* are subject to transcriptional regulation under the *in vitro* conditions we have evaluated.

## Discussion

Bacterial pathogens vary widely in physiological dependence on the host for nutritional support, regulator co-factors, and immune evasion [84–86]. Specifically, trace metals are essential to bacterial growth and virulence, but too little or too much can be problematic. Bacterial metal homeostasis requires tight regulation of metal sensing, transport for accumulation, and efflux to avoid toxicity [30,31,42,85,87]. It is not well understood how *B. burgdorferi* is able to transport necessary nutrients for survival, particularly metals, across the membrane. This is despite the importance of trace metals, such as Zn and Mn, for *B. burgdorferi* regulation being well documented and its uniqueness in lacking a requirement for Fe [18,25,26,29,76,77,79]. Prior studies characterizing borrelial proteins that engage metals have been limited and include a metalloregulatory protein, BosR, and borrelial membrane transporter, BmtA [18,25–27,41,70,76,77,79,88,89]. A previous Tn-seq study identified an annotated Ca^2+^/Na^+^ antiporter *bb0164,* transposon mutant designated as Tn::*bb0164*, as one of many borrelial genes that promote resistance to oxidative stress [20,45,90]. Metal transport was also evaluated in Tn::*bb0164* because *bb0164* was predicted to be an inner membrane transporter with metal binding motifs [20,45]. The Tn::*bb0164* demonstrated a significant reduction in Mn transport, therefore we hypothesized that *B. burgdorferi* has adapted the use of BB0164 to transport key metals that support pathogenic mechanisms and survival during mammalian infection.

An important finding of our study is that *B. burgdorferi* has a second Mn transporter that is not homologous to known bacterial Mn transporters, such as MntH, and is essential for pathogenesis [31,91]. Using a genetic approach, we characterized a borrelial non-polar deletion *bb0164* (*bb0164^-^*) and a chromosomal complement (*bb0164^-/+^*) strain to further characterize the contribution to gene regulation, metal transport, and infectivity. The absence of *bb0164* activity modulates the RpoS cascade, causing elevated expression and production. This overexpression and/or overproduction of the RpoS regulon likely contributes to the loss of infectivity observed in *B. burgdorferi bb0164^-^.* Interestingly, *bosR* expression and BosR production is unchanged in the absence of *bb0164*, suggesting the differences in Mn availability alters BosR activity to promote *rpoS* transcription [41,92–94]. It is possible that lower Mn levels allow for increased or more efficient binding of Zn to BosR resulting in higher levels of active protein. This is supported by work by Troxell et al that demonstrated Mn or Zn supplements individually and in combination were able to transcriptionally regulate the RpoS pathway and post-transcriptionally regulate BosR during cultivation of wild type *B. burgdorferi* [18]. Numerous studies have established that BosR requires Zn as a metal co-factor for dimerization and DNA binding, but the mechanism Mn uses to alter BosR production and/or activity has not been identified [27,76,77,94]. This is further complicated by recent findings that BosR binds RNAs, with one study suggesting that BosR does not bind DNA, but rather stabilized *rpoS* transcript at the 5’ UTR [95–98]. Additionally, BosR chaperone activity and interactions with borrelial sRNA was recently established [97,98]. Metals, including Mn, are known to play a regulatory role by binding riboswitches and promoting the formation of sRNA-protein complexes [42,80,91,99,100]. It is likely that Mn influences regulations at various levels in *B. burgdorferi* that are only just beginning to be discovered.

The virulence regulation and infectivity phenotypes associated with both BB0164 and BmtA sheds light on the reliance on Mn for *B. burgdorferi* pathogenesis. It was unexpected that the loss of one of these Mn transporters while the other remained functional resulted in the observed genetic dysregulation and loss of infectivity [18,25]. Like BmtA, BB0164 exclusively imports Mn and no other important trace metals [18,25]. We postulated that the Mn centric nature of *B. burgdorferi* leant to a mutual regulation of *bb0164* and *bmtA* that may promote availability of the important trace metal, but instead the two transporters are independent of one another and are likely not redundant. Prior work has identified *bb0164* as a member of the BosR regulon and repressed by BosR [101]. This is interesting considering *bb0164* expression increases less than one fold under in vitro mammalian like conditions that are known to induce *bosR* and *rpoS*. *bmtA* expression was not elevated in *bb0164^-^* to compensate for the loss of a Mn transporter under RpoS inducing (37°C, 5% CO_2_) or non-inducing (23°C, atmospheric CO_2_) conditions [14,17,38]. This is also in line with RpoS-independent regulation of *bmtA,* but is counter to the reported temperature regulation of *bmtA* during stationary growth phase [18]. Our findings may differ because *B. burgdorferi* was collected at mid-exponential phase with shifts in temperature and CO_2_. Another potential difference that could have altered intracellular Mn levels was the presence of elevated CO_2_ that could be detrimental to Mn transport [102–104]. WT 5A4-NP1 *B. burgdorferi* had higher Mn levels than reported in Ouyang et al that collected cells grown at 37°C, atmospheric CO_2_ for 9 days; therefore, under the conditions of exponential growth it was found that CO_2_ was not detrimental to Mn transport [25]. Prior work showed that exogenous metal content correlated with intracellular content, therefore it could be assumed that metal availability may serve as a regulatory signal in *B. burgdorferi* [18]. The exogenous metal content did not serve as a signal to repress or activate expression of either Mn transporter, but may have influenced protein production, localization, or transport activity. The concentrations used for supplementation represented near physiologic level or a skewed ratio of Mn to Zn while the spirochetes remained in a rich growth medium [18,105]. It is possible that examining metal toxicity would elicit a different regulatory response in *B. burgdorferi* [87,106]. The availability of trace metals is modulated by both the host and pathogenic bacteria that can alter the immune response effectiveness and infectivity [42,85,107]. The effect on and by metals on *B. burgdorferi* infection requires future study. While in ticks, the predominant metal is Fe which is released during the digestion of the tick bloodmeal [108]. Strategically, *B. burgdorferi* has evolved to not require Fe, thus avoiding the associated toxicity, particularly during the tick blood meal [11].

*B. burgdorferi* requires trace metals for motility and growth that is lost in chelated media supplemented with Excyte in place of normal rabbit serum (NRS) [29]. Our data and others demonstrate a significant reduction in growth rate that slowed motility in chelated complete BSK-II that is rescued by Mn or Zn individually [29]. This suggests a potential physiologic role for trace metals in *B. burgdorferi* that was not observed in host adapted *bmtA*^-^ and is likely due to the presence of *bb0164* [25]. To accurately address a physiologic role for Mn transport in *B. burgdorferi*, a double *bmtA bb0164* mutant is needed for testing in the dialysis membrane (DMC) model [25,109]. Another requirement of *bb0164* for survival in the mammalian host may be the role of Mn combating oxidative stress, as our *B. burgdorferi bb0164^-^* strain validated previously published phenotypes using the transposon mutant [45,110]. The absence of metals in growth media also increased the sensitivity of *B. burgdorferi* to oxidative stress that was rescued by Mn and Zn supplementation. *B. burgdorferi* superoxide dismutase (SodA) protects against ROS using Mn as a regulatory signal and co-factor [79,111]. To date, the only protein associated with combating the oxidative stress response that uses Zn as a metal co-factor is BosR [76,77], though it is questioned if BosR serves as a specific oxidative stress regulator or a generalized environmental response regulator [41,76,112–114]. It is interesting that Zn transporters or scavenging mechanisms, such as metallophores, have yet to be identified in *B. burgdorferi* considering Zn supports growth, motility, oxidative stress, and gene regulation. Trace metal levels fluctuate in the body with Zn being more abundant than Mn, particularly in the blood [115–119]. During acquisition or transmission, the spirochete may benefit from the release of trace metals when the blood meal is digested [120]. However, *B. burgdorferi* does not reach high levels of spirochetemia and is found for a short period of time in the mammalian bloodstream, thus this may not have the most influence for borrelial metal homeostasis [121–123]. The most abundant metal reserves in the body are found in tissues, such as the liver, that *B. burgdorferi* is not known to readily colonize and/or cause Lyme associated pathology [85,124–126]. The tug-o-war between *B. burgdorferi* infection and nutritional immunity is a largely under investigated topic that may shed light on tissue tropism of colonization and inflammation.

When *B. burgdorferi* is interacting with the host tissues and environment, it provides the opportunity to acquire trace metals using non-conventional transporters. *Bacillus subtilis* MntH is an example of a well characterized Mn transporter in bacteria, but *B. burgdorferi* does not encode a similar transporter [20,61,127]. Instead, it encodes two non-canonical Mn transporters [20,25]. Our AlphaFold modeling demonstrated a high-level dissimilarity of BB0164 from MntH and BmtA, whereas BB0164 aligns to a high degree of similarity to a Ca^2+^/Cation antiporter from *Methanococcus jannaschii* that supports ion homeostasis [51,53,62]. Additional investigation is needed to characterize the potential antiporter function of BB0164 and if it contributes to cellular homeostasis or protects against pH and/or salt stress [53,62,128,129]. BB0164 may have a dual function of metal and cation transport to support *B. burgdorferi* virulence and physiologic survival.

The characterization of *bb0164* is not without some limitations, primarily being that BB0164 production was not evaluated for cell localization or environmental regulation. Efforts were unsuccessful in generating antibody or a detectable Flag-tagged BB0164 protein, which is often challenging or impossible with inner membrane proteins. Outside the scope of this study is determining the ability of BB0164 to exchange Na in its role in Mn transport or possibly intracellular pH balance [53,62,128]. Another constraint is the requirement to cultivate *B. burgdorferi* in a complex, undefined media that includes normal rabbit serum, thus making it more difficult to control concentrations of nutrients, including trace metals [130]. It may be advantageous to investigate borrelial metal homeostasis using host-adapted *B. burgdorferi* in future studies to gain a better understanding of how *B. burgdorferi* uses metals during mammalian infection. This work, together with the prior *bmtA* studies, demonstrates *B. burgdorferi* reliance on trace metals and highlights the need for further investigation into these unique Mn transporters [18,25,26,28]. Specifically, future work should investigate if BmtA and BB0164 work together and/or independently during the complete pathogenic lifecycle. It also presents the question of how *B. burgdorferi* infection influences nutritional immunity during mammalian infection and if that leads to the inflammatory pathology associated with Lyme disease.

In summary, our work has demonstrated that *B. burgdorferi* encodes a second Mn transporter that operates uniquely from the predicted annotation, BmtA, and other characterized bacterial Mn transporters [20,25,91]. It was unexpected that *B. burgdorferi* would use two non-conventional Mn transporters that contribute to the essential genetic regulation for mammalian adaptation and infection. Interestingly, no zinc transporters have been identified to date despite a prominent role of Zn dependent regulation in *B. burgdorferi* [18,75–78,131]. Metal homeostasis plays a significant role in borrelial pathogenesis by unknown mechanisms that may be important for physiologic survival and immune modulation in addition to the identified contributions to oxidative stress resistance and virulence regulation.

## Materials & Methods

### Bacterial Strains and Growth Conditions

*B. burgdorferi* was grown in Barbour-Stoenner-Kelley II (BSK-II) medium supplemented with 12% rabbit serum (complete BSK-II, pH 7.6) at 37 °C, 5% CO_2_ unless otherwise noted [130]. Divalent-metal chelated BSK-II was created with Chelex-100 (BioRad) as previously described and used for *B. burgdorferi* cultivation at 32 °C, 1% CO_2_ [18]. When appropriate, *B. burgdorferi* was grown with the following antibiotics: 300 µg/mL kanamycin, 50 µg/mL streptomycin, and 100 µg/mL gentamycin. *Escherichia coli* NEB10β (New England Biolabs) strains were grown on Luria-Bertani (LB) agar plates or LB broth at 37°C with the following antibiotics as necessary: 100 µg/mL spectinomycin and 15 µg/mL gentamycin.

*E. coli* and *B. burgdorferi* strains used in this study are listed in Supplemental Table 1; primers used in this study are listed in Supplemental Table 2. To create the *bb0164*^-^ plasmid, 1.5 kb regions upstream and downstream of *bb0164* were PCR amplified with primers *bb0164-*US-F, *bb0164*-US-R and *bb0164*-DS-F-*Xma*I-*Bam*HI, *bb0164*-DS-R. 200 bp upstream of *bb0166* was amplified with primers P_bb0166_-del-F and P_bb0166_-del-R-*Bam*HI-*Xma*I. Amplicons were combined with overlap PCR and cloned into pCR8/GW/TOPO (Invitrogen) to generate pFL307. The P*_flgB_*-*aadA* antibiotic cassette was amplified from pKFSS-1 using primers *aadA*-F-*Bam*HI and *aadA*-R-*Xma*I [69], digested with *Bam*HI and *Xma*I, and ligated into linear pFL307 to create pFL302. Electroporation of *B. burgdorferi* 5A4-NP1 with pFL302 was performed to generate *bb0164*^-^. The *bb0164^-/+^* plasmid was generated by PCR amplifying the *bb0164* ORF and the *bb0166* upstream region with primers *bb0164*-ORF-F, *bb0164*-ORF-R-*Not*I and P*_bb0166_*-com-F-*Sal*I, P*_bb0166_*-com-R. Amplicons were combined with overlap PCR, digested with NotI and SalI, ligated into linear pVA110 to generate pFL504, and electroporated into *bb0164^-^* to generate *bb0164*^-/+^ [132].

### In silico Analyses

Structural models of BB0164, BB0164::Mn^2+^, BmtA, BmtA::Mn^2+^, *B. subtilis* MntH, and MntH::Mn^2+^ were generated using AlphaFold 3 [133]. NCX_Mj structural coordinates were obtained from PDB IDs 5HWY and 5HXR [62]. Structural visualizations and analyses were performed using UCSF ChimeraX 1.10. Root mean square deviation (RMSD) values were calculated from distances between backbone Cα atoms where applicable. pLDDT and iPTM values were determined from AlphaFold-generated model files; structures were colored using bfactor color command in ChimeraX 1.10. Metal-coordination site geometries were validated with CheckMyMetals [63–66].

### Inductively Coupled Plasma Mass Spectrometry (ICP-MS)

Mid-logarithmic phase *B. burgdorferi* strains were harvested and washed as previously described [26,45]. Briefly, pellets of 10^9^-10^10^ cells total were resuspended in 16N nitric acid, incubated at 100 °C for 15 min, then diluted to 6% nitric acid with H_2_O. Intracellular metal content was determined via inductively coupled plasma mass spectrometry (ICP-MS) with the PerkinElmer NexION 300D ICP-MS housed in the Elemental Analysis Lab at the Texas A&M University Department of Chemistry using a 48 component multi-element ICP-MS standard (Solution A, High-Purity Standards) with rhodium as an internal control.

### ROS and RNS Stress Assays

Mid-logarithmic phase *B. burgdorferi* strains were treated with H_2_O_2_, *tert*-butyl hydroperoxide (TBHP), or diethylamine NONOate (DEA/NO) as previously described [45,134]. For ROS assays, cells were washed, resuspended in modified BSK-II (lacking pyruvate, bovine serum albumin, and rabbit serum), treated with 0, 125, and 185 µM H_2_O_2_, or 0, 1.5, and 2.5 mM TBHP, incubated at 37 °C, 5% CO_2_ for 4 h, then diluted 10-fold in BSK-II until untreated WT reached mid-logarithmic phase. For RNS assays, cells were treated with 0, 1.25, or 2.5 mM DEA/NO for 2 h at 37 °C, 5% CO_2_ in complete BSK-II and diluted as described above. Survival was calculated as the ratio of live cells in treated samples compared to untreated for each strain.

*B. burgdorferi* 5A4-NP1 was grown to mid-logarithmic phase in complete BSK-II, chelated complete BSK-II, or chelated complete BSK-II supplemented with 10 µM MnCl_2_, 100 µM ZnSO_4_, or 10 µM MnCl_2_ and 100 µM ZnSO_4_. Cells were harvested, washed, and resuspended in modified BSK-II, chelated modified BSK-II, or chelated modified BSK-II supplemented with 10 µM MnCl_2_ and/or 100 µM ZnSO_4_, and treated with 185 µM H_2_O_2_ [18]. Cultures were incubated at 32 °C, 1% CO_2_ for 2 h, diluted 10-fold in complete BSK-II, and survival quantitated as described above.

### Mouse Infections

Groups of five 6-8-week old female C3H/HeN mice (Charles River Laboratories) were infected intradermally via needle inoculation with 10^5^ cells WT, *bb0164^-^*, and *bb0164^-/+^ B. burgdorferi*. At 14 and 28 days post-infection (dpi), mice were euthanized and inoculation-site skin flank, ear pinnae, inguinal lymph node, heart, bladder, and tibiotarsal joint were aseptically collected in BSK-II. Infection was determined by *B. burgdorferi* tissue outgrowth up to 1-month post-collection.

### RNA isolation, Reverse-Transcriptase PCR (RT-PCR), and Quantitative Reverse-Transcriptase PCR (qRT-PCR)

Total RNA was isolated from *B. burgdorferi* grown at 37 °C, 5% CO_2_ and 23 °C using the hot phenol extraction method previously described [135,136]. Total RNA was extracted from *B. burgdorferi* grown at 32 °C, 1% CO_2_ in complete BSK-II, chelated BSK-II, and chelated BSK-II supplemented with 0.5 µM MnCl_2,_ 10 µM MnCl_2_, 100 µM ZnSO_4_, or 10 µM MnCl_2_ and 100 µM ZnSO_4_ with the Quick RNA Micro-Prep (Zymo Research) per manufacturer’s instructions. cDNA was created using the Superscript III First Strand cDNA Synthesis Kit (Invitrogen), random hexamers, and 500 ng total RNA. Transcriptional copy number of *bosR, rpoS, ospC, dbpA*, and *flaB* were determined using PerfeCTa SYBR Green FastMix (QuantaBio) and CFX384 Real-Time System (BioRad) via the ΔΔC_T_ method. RT-PCR was performed on cDNA from WT cells grown at 37 °C, 5% CO_2_ using primers described in Supplemental Table 1 and DreamTaq Hot Start PCR Master Mix (ThermoFisher).

### Western Analysis

*B. burgdorferi* whole cell lysates were analyzed by 12.5% sodium dodecyl sulfate-polyacrylamide gel electrophoresis (SDS-PAGE) as previously described [137–139]. Protein production was assessed with mouse antibodies against OspC, FlaB, BosR, and RpoS primaries, goat anti-mouse IgG HRP secondary (ThermoFisher), and Western Lightning-Plus ECL chemiluminescent substrate (Revvity) on film. Lysates and blots were obtained in triplicate.

### Animal Ethics and Welfare Statement

Texas A&M University is accredited by the Association for Assessment and Accreditation of Laboratory Animal Care (AAALAC), indicating a commitment to responsible animal care and use. All animal experiments were performed in accordance with the Guide for Care and Use of Laboratory Animals provided by the National Institutes of Health (NIH) and the Guidelines for the Approval for Animal Procedures provided by the Institutional Animal Care and Use Committee (IACUC) at Texas A&M University.

### Statistical Analyses

All statistical analyses were performed using GraphPad Prism (GraphPad Software, Inc.). Statistical significance was determined by one-way ANOVA tests among three biological and three technical replicates unless otherwise noted. A *p*-value of less than 0.05 was considered statistically significant for all analyses performed.

## Acknowledgements

We thank Dr. Brandon Garcia, Dr. Jon T. Skare, and members of the Skare lab for their valuable insight, helpful discussions, and feedback in the development of this manuscript. We also acknowledge Dr. Bryan E. Tomlin and Kevin LeDone at the Elemental Analysis Laboratory at the Texas A&M University Department of Chemistry’s Center for Chemical Characterization and Analysis for running our processed samples on the ICP-MS and providing valuable insight into the ICP-MS process.

## Supplemental data

**Fig S1.**
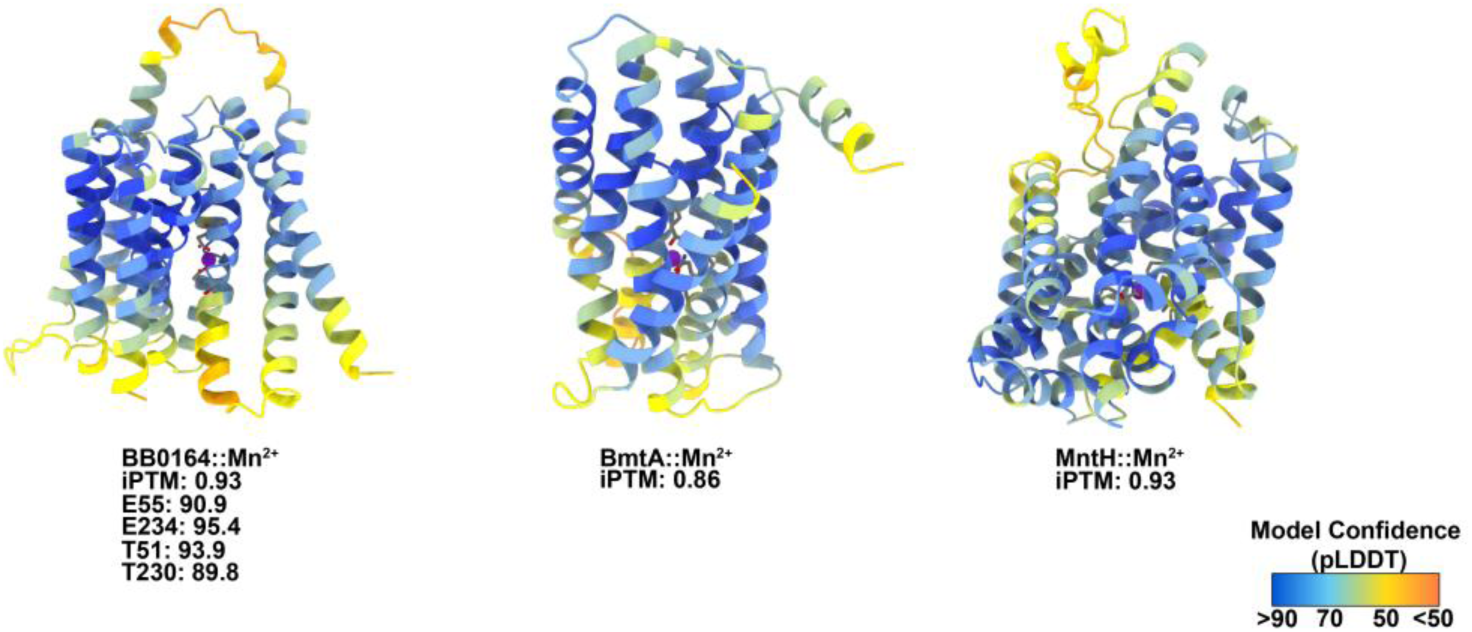
Confidence scores for AlphaFold predicted models. AlphaFold models for cation bound BB0164, BmtA, and MntH were colored according to confidence scores (pLDDT). Lower confidence is seen in N- and C-terminal ends as well as extracellular loops with higher confidence predicted for internal helices. iPTM scores for BB0164 also showed high confidence in the cation-coordinating residues.

**Video submitted in separate file.**

**Fig S2. BB0164 is predicted to undergo conformational shifts when bound to Mn^2+^.** Morph video depicting AlphaFold 3-generated structures of apo BB0164 (green) and BB0164::Mn (tan). The Mn^2+^ ion is depicted as a purple sphere. Helices 1, 2a, 6, and 7a shift across the body of the protein upon Mn^2+^ binding.

**Fig S3.**
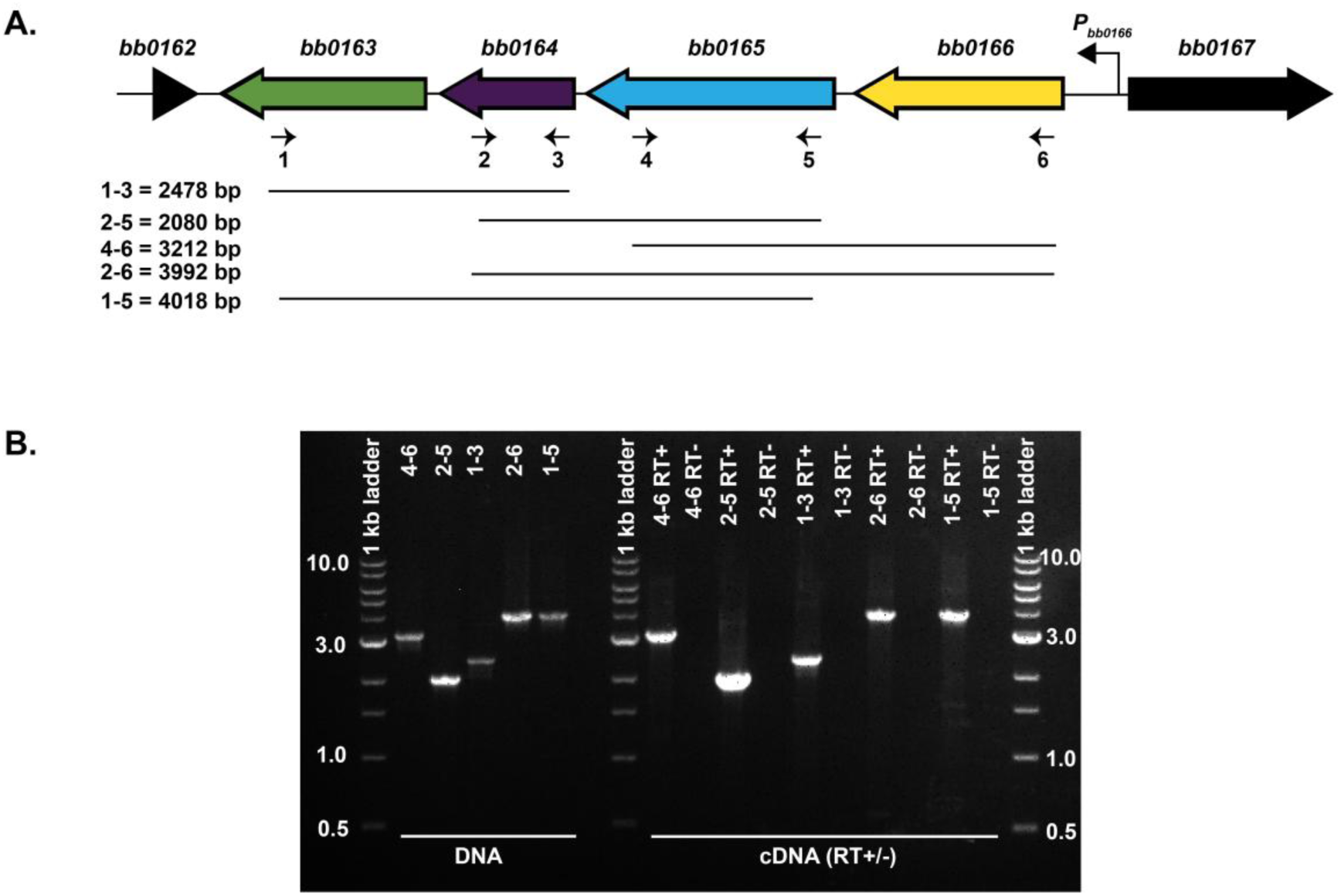
Operon determination for *bb0164*. **A.** Schematic of operon for BB0164 with surrounding genes labelled on large arrows indicating transcription direction. Numbered arrows indicate primer locations. Lines below and base pair numbers represent predicted amplicon length. **B.** 1% TAE gel electrophoresis of indicated gene specific primer sets with *B. burgdorferi* 5A4-NP1 cDNA. DNA reactions shown as positive controls and RT+ and RT- are in sequential lanes.

**Fig S4.**
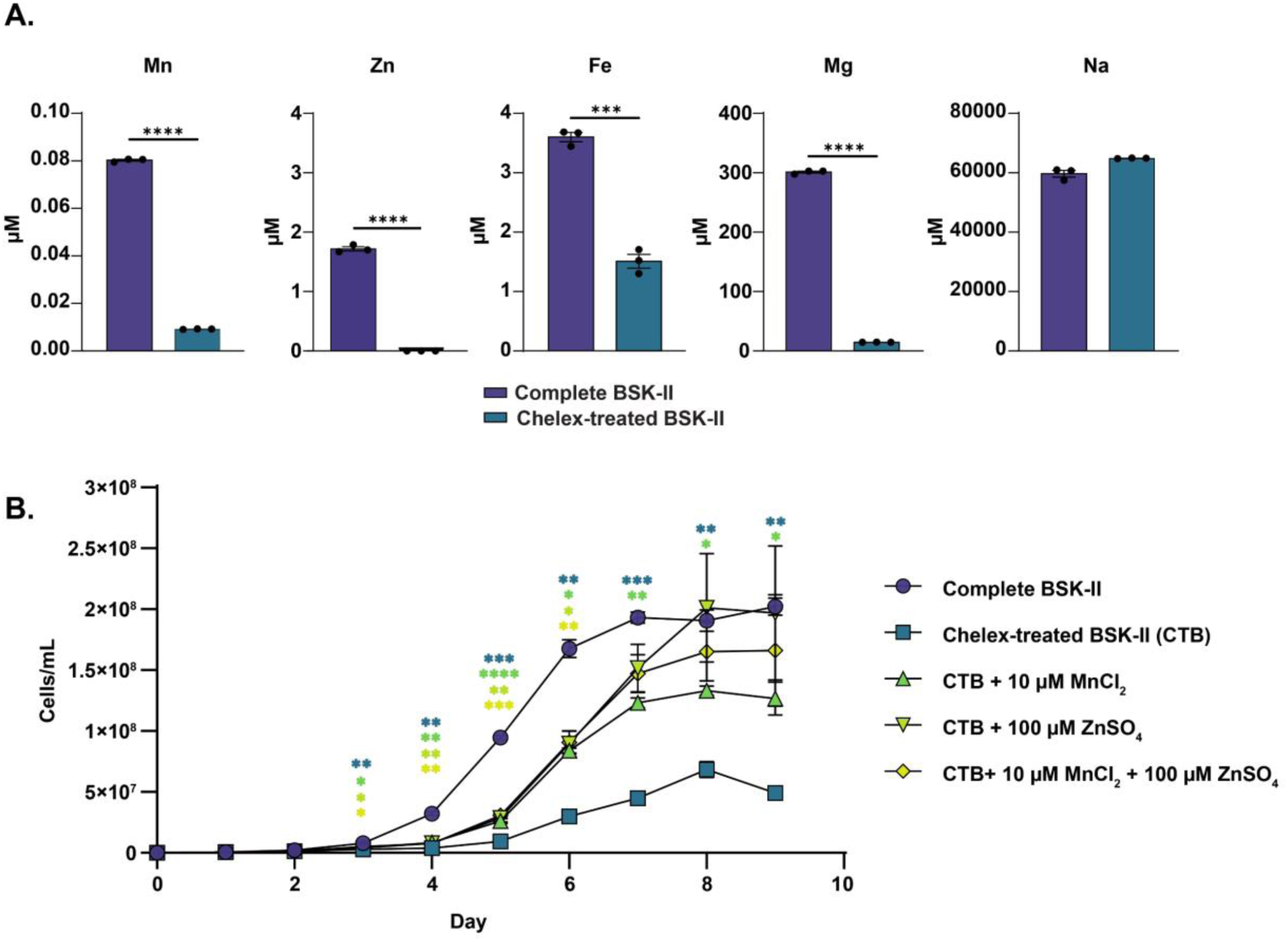
Chelation of divalent metals from BSK-II impedes optimal *B. burgdorferi* growth *in vitro*. **A.** Inductively-coupled plasma mass spectrometry (ICP-MS) measurements of Mn, Zn, Fe, Mg, and Na concentration in normal BSK-II and chelated BSK-II. **B.** Growth curve of WT *B. burgdorferi* grown at 32°C, 1% CO_2_ in normal BSK-II, chelated BSK-II, and chelated BSK-II supplemented with either 10 μM MnCl_2_, 100 μM ZnSO_4_, or a combination of 10 μM MnCl_2_ and 100 μM ZnSO_4_. * p<0.05. ** p<0.01. *** p<0.001. **** p<0.0001.

**Supplemental Table 1:** Strains and plasmids used in this study.

| Strain | Description | References |
| --- | --- | --- |
| ***B. burgdorferi* strains** |  |  |
| 5A4-NP1 (WT) | Clonal infectious isolate of 5A4bbe02 disrupted by $P_{flgB}$-*kanR*, lacking cp9 | [140] |
| *bb0164*<sup>-</sup> | 5A4-NP1::Δ*bb0164-aadA* | This study |
| *bb0164*<sup>-/+</sup> | 5A4-NP1::Δ*bb0164-aadA*::*bb0164-gentR* | This study |
| ***E. coli* strains** |  |  |
| NEB10β | $\Delta$(*ara-leu*) 7697 *araD*139 *fhuA* Δ*lacX*74 *galK*16 *galE*15 *e*14- ϕ80*dlacZ*ΔM15 *recA*1 *relA*1 *endA*1 *nupG* *rpsL* (*Str*<sup>R</sup>) *rph* *spoT*1 $\Delta$(*mrr-hsdRMS-mcrBC*) | New England Biolabs |
| **Plasmids** |  |  |
| pCR8 | Intermediate for TOPO cloning, *specR* | Thermo Fisher |
| pKFSS-1 | Shuttle vector, $P_{flgB}$-*aadA* | [69] |
| pFL307 | pCR8::upstream and *Pbb0166*-downstream regions of *bb0164* | This study |
| pFL302 | pFL307::$P_{flgB}$-*aadA* | This study |
| pVA110 | Allelic exchange vector for insertion into borrelial | [132] |
|  | chromosome intergenic space between *bb0445* and *bb0446*, *specR* and *gentR* |  |
| pFL504 | pVA110:: P<sub>*bb0166*</sub>-*bb0164*, *gentR* | This study |

**Supplemental Table 2:**
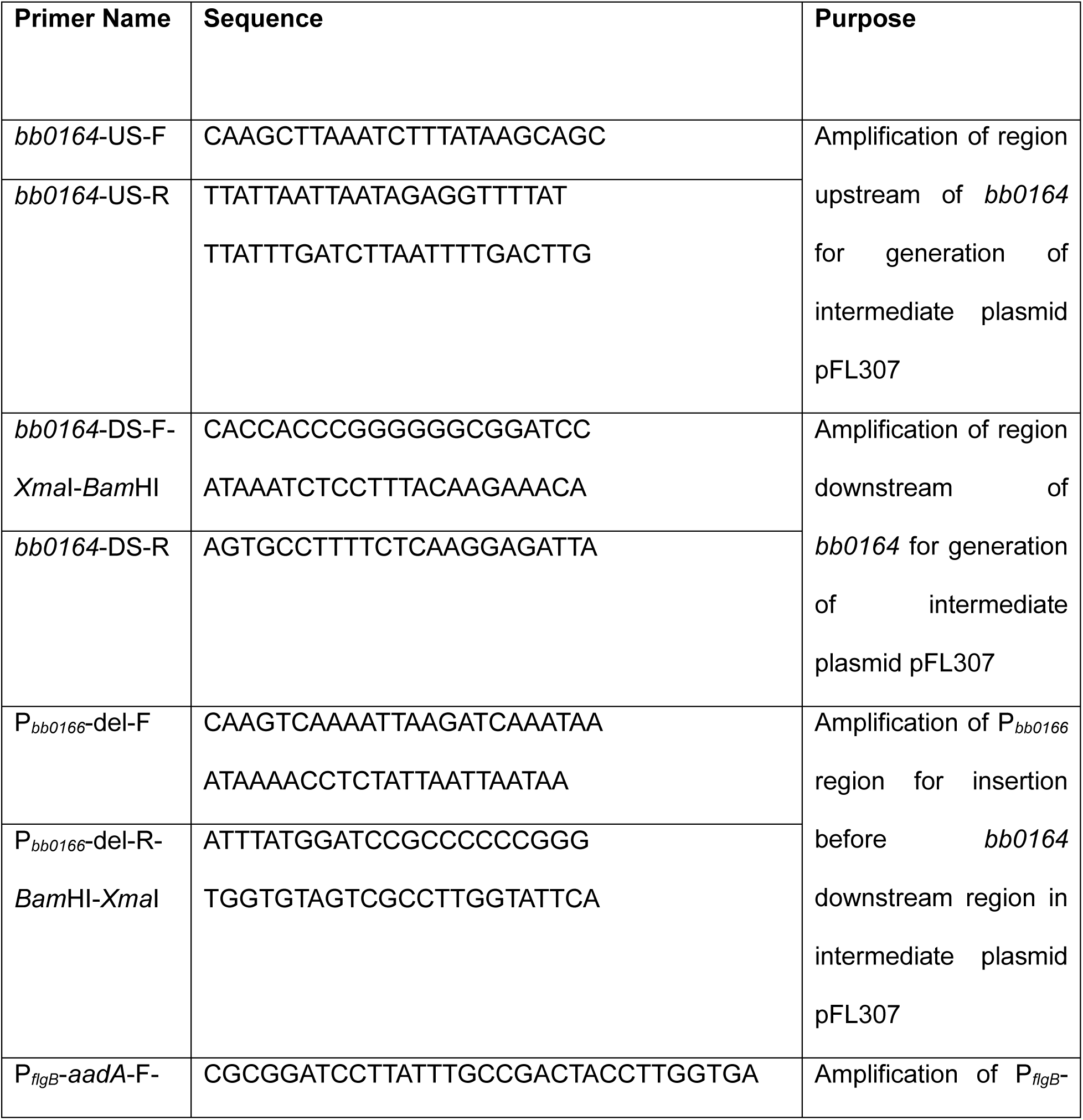
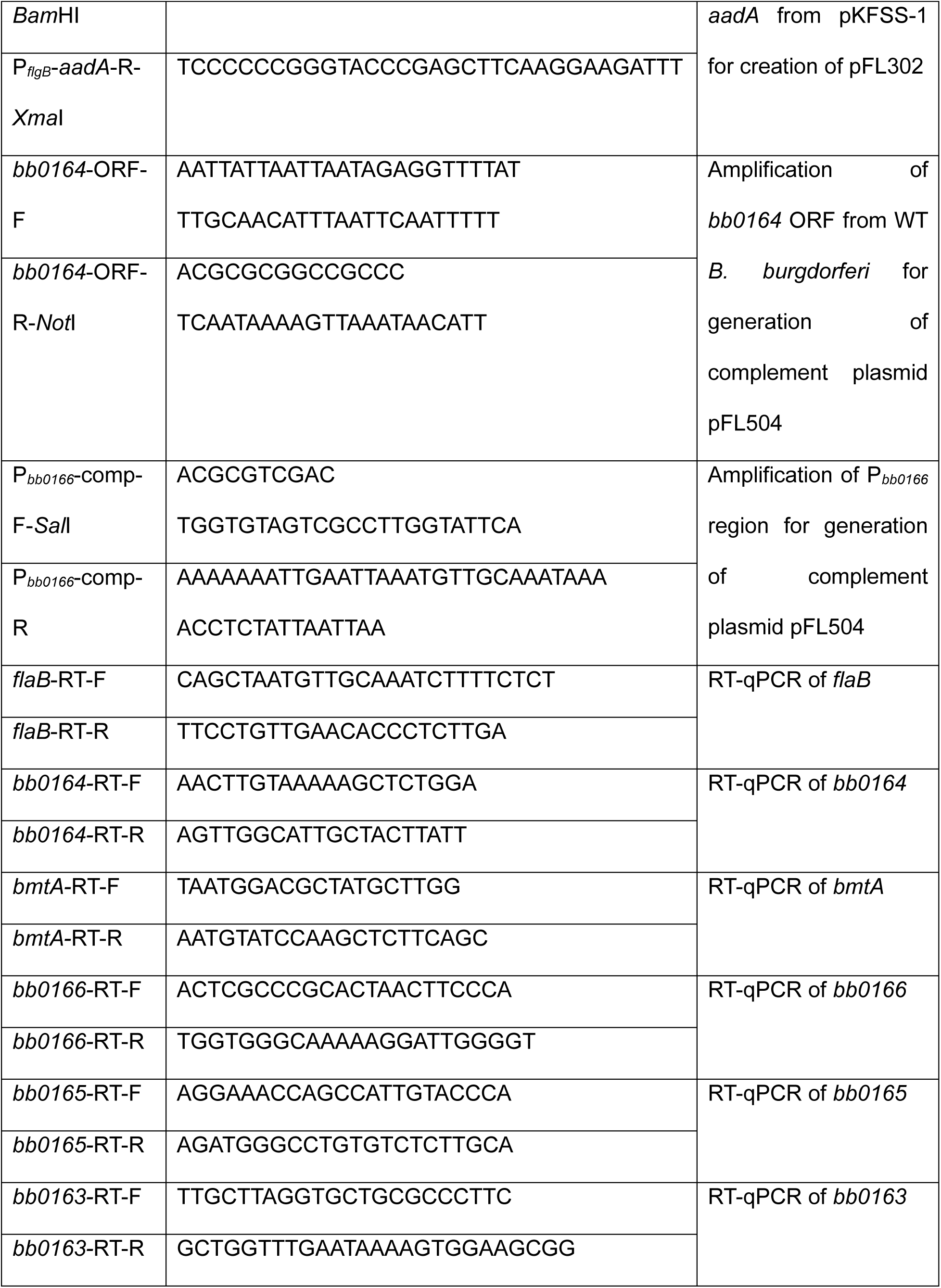
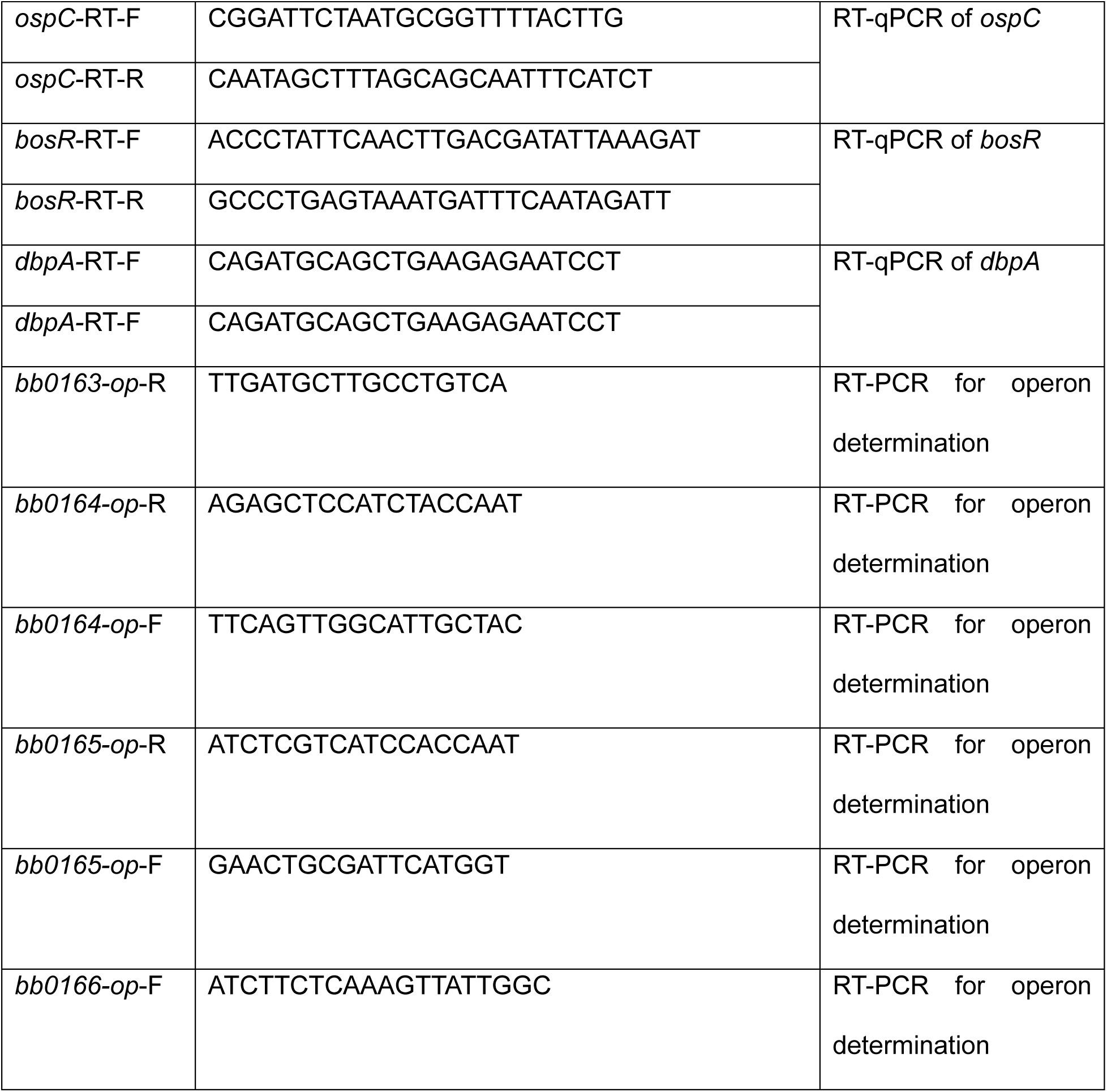
Primers used in this study.

